# Integrating Quality of Life and Survival Outcomes Cardiovascular Clinical Trials: Results from the PARTNER Trial

**DOI:** 10.1101/513895

**Authors:** Jacob V. Spertus, Laura A. Hatfield, David J. Cohen, Suzanne V. Arnold, Martin Ho, Philip G. Jones, Martin Leon, Bram Zuckerman, John A. Spertus

## Abstract

**Background:** Survival and health status (e.g., symptoms and quality of life) are key outcomes in clinical trials of heart failure treatment. However, health status can only be recorded on survivors, potentially biasing treatment effect estimates when there is differential survival across treatment groups. Joint modeling of survival and health status can address this bias.

**Methods and Results:** We analyzed patient-level data from the PARTNER 1B trial of transcatheter aortic valve replacement (TAVR) versus standard care. Health status was quantified with the Kansas City Cardiomyopathy Questionnaire (KCCQ) at randomization, 1, 6, and 12 months. We compared hazard ratios for survival and mean differences in KCCQ scores at 12 months using several models: the original growth curve model for KCCQ scores (ignoring death), separate Bayesian models for survival and KCCQ scores, and a Bayesian joint longitudinal-survival model fit to either 12 or 30 months of survival follow-up. The benefit of TAVR on 12-month KCCQ scores was greatest in the joint model fit to all survival data (mean difference = 33.7 points; 95% CrI: 24.2, 42.4), followed by the joint model fit to 12 months of survival follow-up (32.3 points; 95% CrI: 22.5, 41.5), a Bayesian model without integrating death (30.4 points; 95% CrI: 21.4, 39.3), and the original growth curve model (26.0 points; 95% CI: 18.7, 33.3). At 12 months, the survival benefit of TAVR was also greater in the joint model (hazard ratio = 0.50; 95% CrI: 0.32, 0.73) than in the non-joint Bayesian model (0.54; 95% CrI: 0.37, 0.75) or the original Kaplan-Meier estimate (0.55; 95% CI: 0.40, 0.74).

**Conclusions:** In patients with severe symptomatic aortic stenosis and prohibitive surgical risk, the estimated benefits of TAVR on survival and health status compared with standard care were greater in joint Bayesian models than other approaches.

## BACKGROUND

Patients with heart disease seek treatments to extend survival and optimize their health status; their symptoms, function, and quality of life.^1, 2^ Accordingly, many clinical trials include patient-reported outcomes alongside mortality to measure the impact of treatment on patients’ health status. Traditionally, most trials have reported mortality and health status results independently, such that the health status outcomes are reported only in surviving patients who are able to complete the health status questionnaires. Although this approach has appeal due to its simplicity and ease of interpretation, independently analyzing outcomes may lead to incorrect inferences if the 2 outcomes are inherently related. For example, if sicker patients are more likely to die, then the remaining patients’ average health status is higher than it would be had all patients survived. In studies where a treatment leads to improved survival and health status, ignoring these deaths when analyzing health status outcomes may bias the estimates of health status outcomes, had all patients survived,^3–7^ due to the informative missingness whereby the sickest patients are systematically missing due to censoring by death. Given the correlation between health status and survival, modeling both outcomes simultaneously may improve the precision of both effect estimates^8^ and providing a more holistic summary of treatment benefits. Such an approach could also help integrate these different outcomes when treatment improves health status, but worsens survival, or vice versa.

While a number of potential strategies have been proposed for this purpose,^9–12^ there has been a growing interest in the use of Bayesian models to jointly model survival and longitudinal health status outcomes.^8, 13–16^ A Bayesian approach can enable survival data to inform estimates of longitudinal health status trajectories, thereby partially addressing the problem of informatively missing health status data by estimating an average health status trajectory had all subjects survived. To date, these approaches have mostly focused on improving the power of clinical trials by incorporating patients’ health status into the survival estimates, as opposed to using the survival data to improve the health status estimates.^8, 13^ To clarify the impact of such approaches on the estimates of health status, we applied joint modeling to a cardiovascular clinical trial in which the experimental treatment had an impact on both survival and health status. Specifically, we applied a Bayesian joint model of survival and health status to describe the outcomes of patients with severe, symptomatic aortic stenosis and extreme surgical risk who were not candidates for surgery and were randomized to either transcatheter aortic valve replacement (TAVR) or standard therapy (including a combination of medical management and balloon valvuloplasty) in the Placement of Aortic Transcatheter Valves PARTNER 1B trial.^17, 18^ We compared these estimates with alternative modeling approaches including the original analysis and a more standard Bayesian model that didn’t jointly model the outcomes. These analyses can highlight the potential benefits of a joint-modeling analytic approach using a Bayesian framework to provide a deeper understanding of the benefits of TAVR on patients’ health status and survival.

## METHODS

### PARTNER 1B Trial Data

The PARTNER 1B trial was a randomized, open-label trial designed to test TAVR in patients with severe, symptomatic aortic stenosis for whom surgical aortic valve replacement (SAVR) was considered to present a prohibitive risk for surgery. Eligible patients were randomized to TAVR (performed by a transfemoral approach using the balloon expandable Sapien valve) or standard care, consisting of drug therapy and/or balloon valvuloplasty. As previously described, the primary results of the PARTNER 1B trial demonstrated that TAVR led to substantial improvements in both survival^17^ and health status.^18^ We performed secondary analyses of the patient-level data from this study to examine the impact of a Bayesian joint modeling approach on both outcomes. The study was approved by IRB at each site, and all patients provided informed consent. The Saint Luke’s Health System’s Institutional Review Board reviewed the project and determined this secondary analysis not to be human subjects research.

### Outcomes

Health status was evaluated using the Kansas City Cardiomyopathy Questionnaire (KCCQ),^19^ a 23-item, disease-specific questionnaire that quantifies patients’ symptoms, physical and social limitations, self-efficacy, and quality of life due to heart failure.^19–22^ The KCCQ has undergone extensive reliability and validity testing in various heart failure populations,^19^ including those with severe aortic stenosis.^20^ Each domain is converted to a range of 0-100, where higher scores indicate fewer symptoms, less functional limitation, and better quality of life. The Symptom, Physical Limitation, Social Limitation and Quality of Life scales can be combined to form the KCCQ overall summary scale (KCCQ-OS), which also ranges from 0 to 100, with higher scores indicating better health status. Lower KCCQ scores have been associated with increased risk of death, including in patients undergoing TAVR.^6, 23^ Changes of 5, 10, and 20 points correspond to small, moderate or large clinical improvements, respectively.^21^ The KCCQ was administered prior to randomization and at 1, 6 and 12 months later, providing a longitudinal trajectory summarizing surviving patients’ health status over time. Mortality was measured as death from any cause. At the time of database lock for the quality of life study, all patients had been followed for survival for at least 1 year and up to 30 months.

### Joint Modeling Approach

To address informatively missing health status data due to death, we specified a piecewise linear model for the mean health status trajectories with intercepts at baseline and 1 month and with a linear slope between months 1 and 12. This model (fully specified in Appendix A) was chosen based on observing a large improvement in health status immediately after TAVR or standard therapy, and an approximately linear trend afterwards. Treatment effects were parameterized with two coefficients, one for the difference between treatment and standard care at 1 month and a second for the difference in post-1-month slopes. Baseline average scores were constrained to be the same in the two treatment groups, which was felt to be reasonable owing to randomization and was empirically verified in the study sample. The errors were assumed to be Gaussian and individuals had 3 Gaussian random effects: a baseline intercept, a 1-month intercept, and a post-1-month slope.

Survival times were modeled using a Weibull distribution, with a single shape parameter and a regression model for the scale parameter. The regression model included an indicator of treatment group and two of the individual-level random effects: the 1-month intercept and the post-1-month slope of KCCQ scores. The coefficient on the treatment indicator quantifies the treatment effect in the survival sub-model. The coefficients on the individual-level random effects quantify the relationship between the health status trajectories and hazard of death. Positive coefficients on the random effects would indicate that higher health status scores at 1 month and more positive slopes are associated with shorter survival. In this joint model, survival times also influence the individual-level random effects, which helps account for censoring of health status by death. We specified weakly informative priors following best practices in Bayesian data analysis.^24^

To fit the model to the KCCQ-OS data, we first transformed the scores to make them suitable for a Gaussian model. We trimmed scores of 0 and 100 to 1 and 99, then divided by 100 and applied a probit transformation. Given that the KCCQ-OS does not have a score of 1 or 99, the trimmed scores remained the lowest and highest possible. The resulting transformed KCCQ-OS scores were approximately normally distributed (see Figure S1 in Appendix B). All data analyses were done using R software, version 3.3.1. Model code was written in Stan for Hamiltonian Monte Carlo (HMC) sampling, and fit with the ‘rstan’ package and is provided in Appendix C.^25^ The modeling of means using frequentist t-tests and regression estimates generate confidence intervals (CI) around the point estimates, whereas the Bayesian analyses generate credible intervals (CrI). We assessed convergence of our HMC chains using R-hat statistics, where an R-hat below 1.1 indicated reasonably good convergence, and by visually examining trace plots.^24^

We compared multiple approaches to illuminate features of the analyses that influence the estimates of health status outcomes. These comparators include the original estimates of health status outcomes, which were based on longitudinal growth curve models that ignored death,^18^ and the Bayesian model of health status data we specified with no sharing of information through joint random effects (termed the ‘separate’ Bayesian analysis).

Furthermore, we included two implementations of our joint Bayesian model: one using survival follow up only through to 12 months to match the primary survival analysis^17^ and one using up to 30 months of survival follow up to examine the effect of using all available survival data on joint model fitting and inference. To determine the extent to which survival data actually informed health status estimates in our joint modeling procedure, we examined the survival coefficients of the random 1-month intercept and post-1-month slope random effects. As they are complicated and mostly of technical interest, the results for the joint random effects are presented in Appendix D.

## RESULTS

### Patient Population

Table 1 displays baseline patient characteristics and observed outcomes of the 358 randomized patients. The population was elderly, with a mean age of 83.2 ± 8.4 years, 46.4% were male, and 91.3% were white. By design, the population was at high risk for SAVR (mean STS score of 11.5), and patients had high rates of cerebrovascular disease and chronic obstructive pulmonary disease.

**Table 1:** Baseline Characteristics and 1-Year Outcomes

| Variable | TAVR<br>(N = 179) | Medical Therapy<br>(N = 179) |
| --- | --- | --- |
| Age, years ± SD | 83.1 ± 8.6 | 83.2 ± 8.3 |
| Male sex, % | 45.8 | 46.9 |
| White race, % | 92.2 | 90.5 |
| COPD, % | 41.3 | 52.5 |
| Cerebrovascular Disease, % | 27.4 | 27.4 |
| STS Mortality Risk Score ± SD | 11.2 ± 5.8 | 11.9 ± 4.8 |
| KCCQ-OS |  |  |
| Baseline ± SD | 36.2 ± 20.5 | 34.4 ± 20.1 |
| 1-Month ± SD | 61.6 ± 26.2 | 49.2 ± 24.3 |
| 6-Month ± SD | 70.7 ± 23.0 | 50.5 ± 26.1 |
| 12-Month ± SD | 69.4 ± 25.4 | 47.0 ± 24.6 |
| 1-Year Survival (95% CI) | 0.69 (0.63, 0.76) | 0.49 (0.42, 0.57) |

*SD = standard deviation; STS = Society of Thoracic Surgeons; COPD = chronic obstructive pulmonary disease; CI = confidence interval.*

At 1 month, the raw data indicated significantly more mean KCCQ-OS improvement in the TAVR group than in the standard care group. People in the standard care group had worse KCCQ-OS scores at 12 months than at 1 month, indicating a decline in health status over time. In contrast, patients treated with TAVR experienced improved health status, on average, throughout the year of observation. 1-year survival was also significantly worse in the standard care group than in the TAVR group (49% vs 69%; p<0.01).

### Impact of Joint Modeling on Health Status Inference

Figures 1 and 2 plot the KCCQ-OS results in two distinct ways. Figure 1 plots the estimates from different methods overlaid on each other to facilitate comparison, with separate panels for treatment group. Figure 2 plots each method in a separate panel and includes 95% CIs or CrIs for each estimated trajectory. These plots show that both joint approaches estimated lower health status in the standard care group than either the longitudinal growth curves or the separate Bayesian model. Furthermore, all 3 Bayesian models estimated slightly higher health status in the TAVR group at 12-months, though as we accounted for increasing amounts of survival data the health status trajectories in TAVR-treated patients slightly decreased. Including survival data in the joint model changed the estimated trajectories for the standard care group: the estimate for standard care 12-month KCCQ-OS was 33 from the full survival joint model compared with 44 from the separate Bayesian model and 41 from the longitudinal growth curve model. In terms of inference, the joint methods exhibited wider credible intervals than either the growth curves (which estimated frequentist confidence intervals) or the separate Bayesian model. However, including more survival data led to tighter credible intervals, as shown in the comparison of plots 2C and 2D.

**Figure 1:**
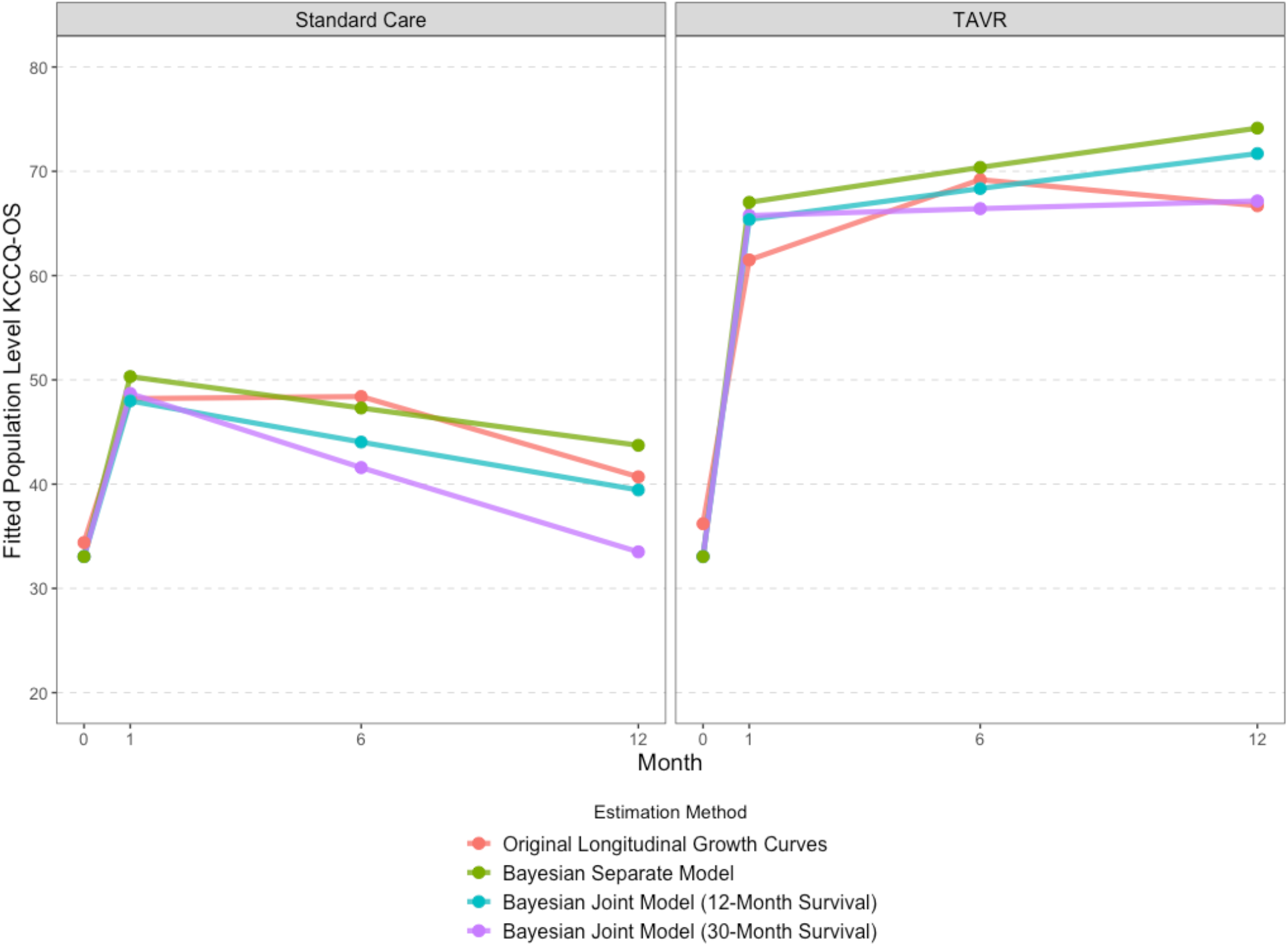
Estimated KCCQ-OS score trajectories for TAVR and standard care according to the original longitudinal growth curves and Bayesian models. Estimated population KCCQ-OS trajectories. Different estimation methods are overlaid and confidence/credible interval ribbons are suppressed for clarity.

**Figure 2:**
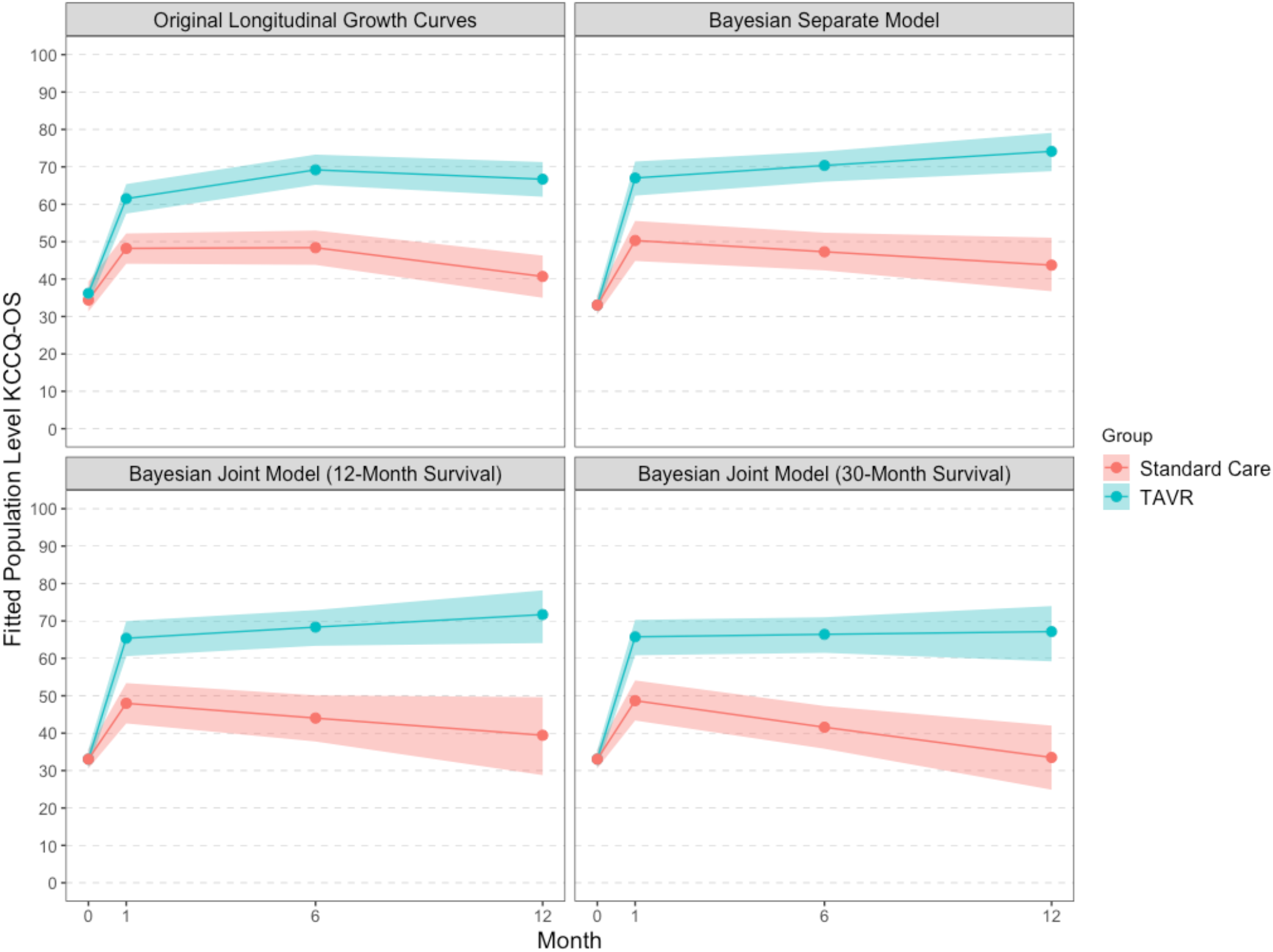
Estimated KCCQ-OS scores for each estimation method with confidence/credible interval ribbons. Estimated population level KCCQ-OS scores from baseline (0-months) to 12-month. Panels show (from left to right, top to bottom) longitudinal growth curve model, Bayesian piecewise linear model fit with no joint parameters, Bayesian piecewise linear model with joint random effects utilizing 12-months of survival data, and Bayesian piecewise linear model with joint random effects utilizing the full 30-months of survival data. Solid lines with points are joint model estimates, ribbons are 95% credible intervals. Standard care group is in red, TAVR is blue.

These differences were also apparent in the effect estimates, defined as the difference in estimated KCCQ scores between TAVR and standard therapy health status at each timepoint (Table 2). The joint Bayesian methods estimated a larger treatment benefit than either the original growth curve model or the separate Bayesian model. In addition, using the full 30-months of survival follow-up data led to a slightly larger KCCQ effect estimate than using only 12 months of survival data.

**Table 2:** Estimated Health Status Benefits

|  | Original Longitudinal<br>Growth Curves<br>(95% CI) | Separate<br>Bayesian<br>Model<br>(95% CrI) | Joint Model<br>with 12-Month<br>Survival Data<br>(95% CrI) | Joint Model<br>with 30-Month<br>Survival Data<br>(95% CrI) |
| --- | --- | --- | --- | --- |
| **1-Month KCCQ-OS**<br>**Effect** | 13.3 (7.6, 19.0) | 16.7 (9.6, 23.7) | 17.4 (9.9, 24.4) | 17.1 (9.8, 24.0) |
| **6-Month KCCQ-OS**<br>**Effect** | 20.8 (14.7, 27.0) | 23.1 (16.3,<br>29.7) | 24.3 (17.5,<br>30.9) | 24.8 (18.0,<br>31.5) |
| **12-Month KCCQ-OS**<br>**Effect** | 26.0 (18.7, 33.3) | 30.4 (21.4,<br>39.3) | 32.3 (22.5,<br>41.5) | 33.7 (24.2,<br>42.4) |

*KCCQ-OS Effect = Difference in Estimated Mean KCCQ-OS between the TAVR and standard of care groups, CI = confidence interval, CrI = credible interval.*

### Impact of Joint Modeling on Survival Inferences

The hazard ratio of TAVR is 0.50 (95% CrI: 0.32, 0.73) from the joint model, which indicates a larger estimate of treatment benefit compared with the original finding of a hazard ratio of 0.55 (95% CI: 0.40, 0.74), or compared to a Bayesian Weibull model with no joint parameters (0.54; 95% CrI: 0.37, 0.75).

## DISCUSSION

As patient-centered outcomes assume increasing importance in both clinical trials and shared decision-making, there is a need for analytic strategies to obtain estimates of the health status benefits when there is informatively missing data, such as censoring by death. This is particularly important in conditions where mortality is common, including heart failure and severe aortic stenosis. In this study, we jointly modeled survival and health status using a Bayesian framework to estimate the impact of mortality on the longitudinal health status benefit of TAVR and to refine the estimates of the survival benefit from TAVR. Integrating mortality and health status demonstrated increased treatment benefits of TAVR compared with medical therapy on both outcomes. Thus, the true benefits of TAVR (in terms of both health status and survival) may have been underestimated by the original analyses, which ignored informative censoring and treated each endpoint separately. These findings illustrate the potential value of joint modeling in providing a more accurate estimate of the impact of therapy in trials where there is a substantial mortality difference between treatment groups that contributes to informatively missing health status data.

Our work extends prior research on the use of joint models to analyze clinical trials. To date, joint modeling has been largely restricted to clinical trials in oncology and HIV/AIDS. While this approach may reflect a greater focus in those fields on analyzing treatments in terms of both survival and quality of life, joint modeling has also been used to improve the precision of survival hazard estimates using longitudinal biomarkers or patient-reported outcomes.^8, 13, 15, 16^ In contrast, cardiovascular trials have traditionally focused on survival or other discrete events, and have only recently begun to emphasize health status outcomes, which have been typically analyzed separately and reported “among survivors,” as in the original analyses of PARTNER 1B.^18^ Indeed, to our knowledge, this is the first application of joint modeling of health status and survival data in a cardiovascular clinical trial.

In particular, applying a joint Bayesian model can enable more accurate assessment of treatment benefits among patients with a highly mortal cardiovascular condition. Previous treatment estimates derived from surviving patients describe the average health status of survivors. However, since patients with worse health status are also less likely to survive, this analytic approach may underestimate the true health status benefit of treatment. In contrast, by explicitly considering the interrelationship between survival and health status, the joint modeling approach provides an estimate of the health status benefit that individual patients would be expected to achieve had they survived to 12 months. We believe that this is the type of information that a prospective patient would want to know when trying to decide whether to undergo a major procedure.

In terms of 12-month survival, the joint model assuming a Weibull distribution estimated slightly better survival compared with the Kaplan-Meier estimate, especially in the TAVR group. Indeed, the hazard of death for TAVR treatment was 50% lower than standard therapy, which was slightly larger than the originally reported treatment effect of a 45% reduction in mortality.^17^

### Limitations

This study should be interpreted in the context of several potential limitations. First, given the study design, there were relatively few health status measurements per person (no more than 4). In using 3 parameters (a baseline intercept, a 1-month intercept, and a post 1-month slope) to define each person’s recovery, the model fit many parameters to relatively sparse longitudinal data. In the future, collection of more longitudinal measurements per subject could allow for more flexible models and lead to improved precision. Modeling a bounded health-status outcome was also challenging. We opted to transform the KCCQ-OS scale to fit a Gaussian model and then back-transform for inference. Our Bayesian approach facilitated this, as we could transform posterior draws and still ensure correct inference. However other modeling choices are defensible, and the treatment effect estimates may be sensitive to the parametric form of the model and data transformations. For future studies, alternative approaches, including a Beta or zero-augmented Beta distribution to model the outcome on its original scale, should be considered.^14^ Moreover, we had to make additional assumptions about the data in order to use parametric sub-models for the longitudinal trajectories and survival, e.g. that post-1-month health status trajectories were approximately linear and survival could be modeled as a Weibull. These assumptions were necessary to fit a joint model that could account for differential survival, despite having limited longitudinal health status measurements. If more data points were available, these assumptions could be relaxed by fitting more flexible sub-models. In addition, while the PARTNER 1B trial was an excellent study with which to explore joint modeling methods given the profound influence of TAVR on both survival and health status, the impact of a joint modeling approach in less effective treatments needs to be explored. Finally, these methods remain unfamiliar to clinicians. While we have attempted to provide intuitive explanations for our approach and for the interpretation of our results, this work represents an early application of joint modeling in cardiovascular disease. Greater experience with these methods and alternative explanations may improve the ability to convey the results of analyses that integrate survival and health status.

### Conclusions

In summary, this study provides the first example in the cardiovascular literature of integrating survival and health status outcomes in a randomized clinical trial using a joint Bayesian framework. We found that in patients with severe, symptomatic aortic stenosis and extreme surgical risk who were randomized to either TAVR or standard medical therapy, the estimated benefits of TAVR on both survival and health status were greater than those observed in analyses that considered those outcomes separately. Since death informatively censors health status observations, this approach may provide a better means of integrating these two clinically important outcomes than merely assuming that the observed health status scores are representative of the overall population of patients in a clinical trial. Future studies should consider incorporating these methods to generate a more accurate estimate of the survival and health status benefits of cardiovascular therapies.

