## Supplementary material for "Integrating Quality of Life and Survival Outcomes Cardiovascular Clinical Trials: Results from the PARTNER Trial": Code: joint_model_final.Rmd

````
---
title: "Joint Survival and KCCQOS Modeling of PARTNER Data"
author: "Jacob Spertus"
date: ''
output:
  html_document: default
  #slidy_presentation: default
  #beamer_presentation: default
header-includes:
- \usepackage{amsmath}
- \usepackage{graphicx}
---

```{r setup, include=FALSE}
#rmd for joint modeling project for paper: Integrating Quality of Life and Survival Outcomes Cardiovascular Clinical Trials: Results from the PARTNER Trial
#Code author: Jacob Spertus
#Paper authors: Jacob Spertus, Laura Hatfield, David Cohen, Suzanne Arnold, Martin Ho, Phil Jones, Martin Leon, Bram Zuckerman, John Spertus

#NB: I use "QOL" and "KCCQOS" interchangeably in the following comments and variable names 
knitr::opts_chunk$set(dev = 'png', echo = FALSE, warning = FALSE, message = FALSE, caption = FALSE)
options(scipen = 20)
seed <- 1234
set.seed(seed)
library(survival)
library(survminer)
library(reshape2)
library(purrr)
library(tidyr)
library(dplyr)
library(mice)
library(broom)
library(rstanarm)
library(lubridate)
library(magrittr)
library(rstan)
library(ggplot2)
source("read_partners_data.R")
setwd("~/Dropbox/FDA/R/Joint Model/Final/")
```


```{r configure data, include = TRUE}
#relu function
relu <- function(x){pmax(0,x)}
#function to transform QOL to be unbounded
to_norm <- function(qol){
  qol[qol >= .99] <- .99
  qol[qol <= .01] <- .01
  qnorm(qol)
}
#function to center and scale time to [-.5,.5]
center_time <- function(x){
  (x - 6)/12
}  
#function to generate a single imputation of missing longitudinal data using PMM (this should be improved in later iterations)
impute <- function(data){
  complete(mice(data, m = 1, method = 'pmm'))
}

#these variables must be characters for proper joinings
data_list$patientdatelist$Subject %<>% as.character()
data_list$baseline$Subject %<>% as.character()
data_list$qol_wide_cat$Subject %<>% as.character()


####################### QOL data ###################
#formatting QOL data
qol_long <- data_list %>%
  pluck("qol_long") %>%
  select(Subject, startdate_ITT, FU_MDY_DT, FolderName, trialarm, time, KCCQOS) %>%
  mutate(VisitNum = as.numeric(recode(FolderName, "Baseline Visit" = 0, "30 Day Follow Up" = 1, "6 Month Follow Up" = 2, "1 Year Follow Up" = 3))) %>%
  mutate(FU_MDY_DT = mdy(format(FU_MDY_DT, "%D"))) 

#further formatting of QOL data
#there are 21 missing KCCQOS scores, we do complete case analysis here (they are dropped)
#also do not keep baseline scores (differences in treatment are reflected in intercept)
qol_data <- qol_long %>% 
  dplyr::select(Subject, time, trialarm, KCCQOS) %>%
  mutate(years = time / 12) %>%
  mutate(KCCQOS = KCCQOS / 100) %>% #rescaled KCCQOS to [0,1] 
  na.omit() %>%
  mutate(NormalKCCQOS = to_norm(KCCQOS)) %>%
  mutate(treat_group = ifelse(trialarm == "TEST", "Treatment", "Control")) %>%
  arrange(Subject, years) %>%
  as_tibble() 

############ survival data ##############
#For censoring at 1 year survival, I use the prefix "Capped"
survival_data <- data_list[c('patientdatelist','baseline','qol_wide_cat')] %>%
  reduce(left_join, by = "Subject") %>%
  dplyr::select(Subject, DaysForITTanalysis, DeathDate, TrialArm, female, racegroup, age, bmi, height, weight, nyha, EF, STS_RISK_SCORE, copd_combined, CV_COND_CEREBRO_YN_foranalysis) %>%
  mutate(gender = ifelse(female == 1, "female", "male")) %>%
  mutate(TrialArm = ifelse(TrialArm == "TEST", 1, 0)) %>% 
  mutate(YearsForITT = (DaysForITTanalysis + 1) / 365) %>%
  mutate(Censored = is.na(DeathDate)) %>%
  mutate(CappedDeathDate = ifelse(YearsForITT > 1, NA, DeathDate)) %>% #for cens indicator
  mutate(CappedCensored = is.na(CappedDeathDate)) %>% 
  mutate(CappedYearsForITT = pmin(YearsForITT, 1)) %>% #truncate censor date to 1 yrs
  mutate(CappedDaysForITTanalysis = CappedYearsForITT*365)

#getting covariate information
mean_sd_age <- survival_data %>% summarize(mean = mean(age), sd = sd(age))
males <- survival_data %>% summarize(pct = mean(gender == "male"))
white <- survival_data %>% summarize(pct = mean(racegroup == "Caucasian"))

#build table 1 split out by treatment group (TrialArm)
table_1_covariates <- survival_data %>%
  group_by(TrialArm) %>%
  summarize(pct_male = mean(gender == "male"), pct_white = mean(racegroup == "Caucasian"), mean_age = mean(age), sd_age = sd(age), mean_sts = mean(STS_RISK_SCORE, na.rm = T), sd_sts = sd(STS_RISK_SCORE, na.rm = T), pct_copd = mean(copd_combined, na.rm = T), pct_cvdisease = mean(CV_COND_CEREBRO_YN_foranalysis, na.rm = T))


################# survival and qol data matching ##################
#the below matches the survival and qol patients. 
#Alignment is important for passing on to Stan joint model.
#Match qol and survival data at the beginning of both lists.
#survival may have more subjects because there might be some subjects without any qol data
#we organize the survival data so all subjects with both QOL and survival appear at start
#additional subjects (those who died and have no QOL data) are indexed at the end of survival

#find index that would give correct sorting of survival to match order of QOL data
qol_survival_index <- match(unique(qol_data$Subject), survival_data$Subject)
#identify subjects in survival who aren't in QOL
no_qol_subjects <- survival_data$Subject[!(survival_data$Subject %in% qol_data$Subject)]
#paste together index for survival subjects who are in QOL and index for survival subjects who are NOT in QOL
final_survival_index <- c(qol_survival_index, match(no_qol_subjects, survival_data$Subject))
#use the survival index to correctly order survival data
final_survival_data <- survival_data %>%
  slice(final_survival_index)
```


# PARTNER Data

- Transcatheter aortic valve replacement (TAVR) in patients not eligible for surgical valve replacement
- $n_1 = 179$ were randomized to TAVR and $n_0 = 179$ were randomized to standard therapy
- Quality-of-life (QOL) data collected using Kansas City Cardiomyopathy Questionnaire (KCCQ) at 0, 1, 6, and 12 months
- Survival data was also recorded, and all subjects were followed for at least 1-year after implantation
- Initial analyses showed substantially improved health status and reduction in mortality due to treatment (Reynolds et al 2011, Leon et al 2010). 
- 1-yr hazard ratio: 0.55 (95\% CI, 0.40-0.74)
- 6-month KCCQ summary difference: 21 points (95\% CI, 15-27)


# Table 1

```{r table 1}
#function to nicely concatenate length-3 vector of form c(mean, lower_ci, upper_ci)
vector_to_string <- function(x){paste(x[1], " (", x[2], ", ", x[3], ")", sep = "")}

#survival analyses should be based on 1 year survival for comparability with original NEJM paper
observed_survival_data <- data.frame(Y = final_survival_data$CappedYearsForITT, Cen = 1-as.numeric(final_survival_data$CappedCensored), Treatment = ifelse(final_survival_data$TrialArm == 1, "Treatment", "Control"))

group_size <- observed_survival_data %>% 
  group_by(Treatment) %>% 
  summarize(Count = n()) %>% 
  spread(key = Treatment, value = Count)


#1-year survival summaries
surv_object <- Surv(time = observed_survival_data$Y, event = observed_survival_data$Cen)
km_treatments <- survfit(surv_object ~ Treatment, data = observed_survival_data)
year_1_survival <- summary(km_treatments, time = 1)
year_1_survival_treatment <- c(year_1_survival$surv[2], year_1_survival$lower[2], year_1_survival$upper[2]) %>% round(2) %>% vector_to_string()
year_1_survival_control <- c(year_1_survival$surv[1], year_1_survival$lower[1], year_1_survival$upper[1]) %>% round(2) %>% vector_to_string()

#in original NEJM paper, 1 year survival is as follows:
#Number at risk: 83 in control, 122 in TAVR
#Mortalities: 89 in control, 55 in TAVR
#In our data:
#Number at risk: 85 in control, 124 in TAVR
#Mortalities: 90 in control, 55 in TAVR
#So basically we have a later version of the dataset that has follow up on more people


#KCCQOS summaries
kccqos_summaries <- qol_data %>%
  group_by(time, trialarm) %>%
  summarise(mean = 100*round(mean(KCCQOS), 3), sd = 100*round(sd(KCCQOS), 3)) %>%
  unite(mean_sd, mean, sd, sep = " +- ") %>% 
  spread(key = trialarm, value = mean_sd)

#make and print table 1
table_1 <- data.frame(
  "Variable" = c("Group Size", "Baseline KCCQOS", "1-Month KCCQOS", "6-Month KCCQOS", "12-Month KCCQOS", "1-year Survival"),
  "Treatment" = c(group_size$Treatment, kccqos_summaries$TEST, year_1_survival_treatment),
  "Control" = c(group_size$Control, kccqos_summaries$CONTROL, year_1_survival_control)
)

knitr::kable(table_1, caption = c("Key outcome variables by treatment group"))
```


# Pre-Processing Steps

- KCCQOS was rescaled to [0,1]. 
- Scores equal to 0 or 1 were trimmed to 0.01 or 0.99 and then KCCQOS was made unbounded by applying the probit transform. 
- After fitting, KCCQOS was transformed back to [0,1].
- Survival time is computed in years. Censoring is recorded if subjects did not die in follow-up period. 
- Baseline measurements are incorporated by using a piecewise linear model. 

# Probit Transformed Data
After transforming KCCQOS data is unbounded and sort of gaussian.

```{r plot transformed data}
#make QOL data long by melting original and transformed KCCQOS into plot
melted_qol_data <- qol_data %>%
  gather(key = "type", value = "score", KCCQOS, NormalKCCQOS) %>%
  mutate(type = ifelse(type == "KCCQOS", "Original KCCQOS", "Transformed KCCQOS"))

transformed_data_plot <- ggplot(melted_qol_data, aes(score)) +
  geom_histogram() +
  facet_grid(. ~ type) +
  xlab("QOL Value")
transformed_data_plot
```

# QQ-Plot of Transformed Data
-We plot the quantiles of our transformed KCCQOS against the quantiles of the normal distribution
```{r plot transformed data qqplot}
#just a different visual of the above
kccqos_qqplot <- ggplot(data = melted_qol_data, aes(sample = score)) +
  facet_grid(. ~ type, scales = "fixed") +
  stat_qq(size = 1.8, alpha = .3) +
  stat_qq_line() + 
  ylab("Sample Quantiles") + 
  xlab("Theoretical Normal Quantiles")

kccqos_qqplot
```


# Joint Model Outline

- Build separate population level models for KCCQOS and for survival
- Simple distributions: KCCQOS is Gaussian (after transform), survival is Weibull  
- Treatment effects: 
  \begin{itemize}
    \item KCCQOS: Treatment determines distinct 1-month KCCQOS and trajectory after 1-month ($\gamma_1$, $\gamma_2$)
    \item Survival: Treatment specific hazard ($\beta_T$)
  \end{itemize}
- Add latent subject-level parameters to KCCQOS model: baseline, 1-month, slope after 1-month ($\theta_{0i}$, $\theta_{1i}$, $\theta_{2i}$)
- Subject-level 1-month KCCQOS parameter is shared with survival model and scaled by $\beta_{\theta_1}$; slope is shared and scaled by $\beta_{\theta_2}$


# Joint Model Specifics

$i \in \{1,...,N\}$ indexes subjects; $j \in \{0,1,6,12\}$ indexes time points. $Y_{ij}$ gives the KCCQOS score at time $j$ for subject $i$; $T_i$ is treatment; $S_i$ is survival time and is possibly right censored. 

 

\begin{align*}
  Y_{ij} &= (\mu_{\theta_0} + \theta_{0i}) \cdot I(j=0) + (\mu_{\theta_1} + \gamma_1 T_i + {\theta_{1i}}) \cdot I(j \geq 1) +\\ &~~~~(\mu_{\theta_2} + \gamma_2 T_i + {\theta_{2i}}) \cdot \max(0, j - 1) + \epsilon_{ij}\\
  S_i &\sim \mbox{Weibull}\{\alpha, \exp(-(\beta_0 + \beta_T \cdot T_i + \beta_{\theta_1} \cdot {\theta_{1i}} + \beta_{\theta_2} \cdot {\theta_{2i}}))\}
\end{align*}


\begin{align*}
  [\mu_{\theta_0}, \mu_{\theta_1}, \mu_{\theta_2}] &\sim \mathcal{N}(0,1)\\
  [\gamma_1, \gamma_2] &\sim \mathcal{N}(0,1)\\
  \epsilon_{ij} &\sim \mathcal{N}(0, \sigma^2)\\
  \theta_{0i} &\sim \mathcal{N}(0, \sigma_{\theta_0}^2)\\
  {\theta_{1i}} &\sim \mathcal{N}(0, \sigma_{\theta_1}^2)\\
  {\theta_{2i}} &\sim \mathcal{N}(0, \sigma_{\theta_2}^2)\\
  %[\beta_T, \beta_0] &\sim \mathcal{N}(0,10)\\
  [\beta_{\theta_1}, \beta_{\theta_2}] &\sim \mathcal{N}(0,10)\\
  [\sigma, \sigma_{\theta_0}, \sigma_{\theta_1}, \sigma_{\theta_2}] &\sim t_3^+(0,1)
  %\alpha &\sim \mathcal{N}^+(0,10)
\end{align*}


- $\mu_{\theta_0}$ = baseline mean; $\theta_{0i}$ = baseline random effect;

- $\mu_{\theta_1}$ = 1-month mean; ${\theta_{1i}}$ = 1-month random effect; $\gamma_1$ = 1-month treatment effect;

- $\mu_{\theta_2}$ = population slope; $\gamma_2$ = treatment specific slope;

- $\epsilon_{ij}$ = error term in QOL model;

- $\beta_0$ = baseline hazard; $\beta_T$ = treatment hazard;

- $\beta_{\theta_1}$ = effect of $\theta_{1i}$s on hazard;

- $\sigma$ = standard deviation of error term; $\sigma_{\theta_0}$ = dispersion of baseline random effect; $\sigma_{\theta_1}$ = dispersion of 1-month random effect;

- $\alpha$ = survival curve shape.


```{r fit model, results = "hide", message = "hide"}
#NB: lines to run stan code are currently commented out, instead saved past runs are loaded for efficiency

#set up data lists that will be passed on to Stan
#Variable names:
#N = total number of observations in QOL data
#I = distinct subjects overall (from (more comprehensive) survival data)
#KCCQ = KCCQ scores (transformed to normality here)
#Timepoint = vector of timepoints for each KCCQOS observation 
#Treatment = vector of treatment group from survival data
#Subjects = vector where each element identifies a distinct subject in the QOL data
#Y = survival time (censored or not)
#Cen = censoring indicator 


############## Spline fit #############
#1 year capped survival data
capped_data_list_spline <- list(
  N = nrow(qol_data),
  I = n_distinct(final_survival_data$Subject),
  KCCQ = qol_data$NormalKCCQOS,
  Timepoint = qol_data$time,
  Treatment = final_survival_data$TrialArm,
  Subjects = as.numeric(as.factor(qol_data$Subject)),
  Y = final_survival_data$CappedYearsForITT,
  Cen = as.numeric(final_survival_data$CappedCensored)
)
#full survival data
data_list_spline <- list(
  N = nrow(qol_data),
  I = n_distinct(final_survival_data$Subject),
  KCCQ = qol_data$NormalKCCQOS,
  Timepoint = qol_data$time,
  Treatment = final_survival_data$TrialArm,
  Subjects = as.numeric(as.factor(qol_data$Subject)),
  Y = final_survival_data$YearsForITT,
  Cen = as.numeric(final_survival_data$Censored)
)
#separate_model_spline <- stan_model("separate_model_spline.stan")
#separate_fit_spline_capped <- sampling(separate_model_spline, data = #capped_data_list_spline, chains = 4, cores = 2, iter = 1000, control = list(adapt_delta = .99, max_treedepth = 14))
#save(capped_separate_fit_spline, file = "separate_fit_spline_capped")
load("capped_separate_fit_spline")


#joint_model_spline <- stan_model("joint_model_spline.stan")
#capped joint fit
#capped_joint_fit_spline <- sampling(joint_model_spline, data = capped_data_list_spline, chains = 4, cores = 2, iter = 1000, control = list(adapt_delta = .99, max_treedepth = 14))
#save(capped_joint_fit_spline, file = "capped_joint_fit_spline")
load("capped_joint_fit_spline")
#joint_fit_spline <- sampling(joint_model_spline, data = data_list_spline, chains = 4, cores = 2, iter = 1000, control = list(adapt_delta = .99, max_treedepth = 14))
#save(joint_fit_spline, file = "joint_fit_spline")
load("joint_fit_spline")
```


```{r get model parameters}
#extract important data from fit
pars_spline <- tidyMCMC(joint_fit_spline, conf.int = TRUE, conf.level = .95) %>% as_tibble()
subjects_frame_spline <- pars_spline %>%
  filter(grepl("bounded_predictor", term)) %>%
  mutate(subjects = data_list_spline$Subjects, time = data_list_spline$Timepoint, observed = qol_data$KCCQOS, treatment = qol_data$trialarm, residual = estimate - observed) 
```


# Observed KCCQOS Trajectories 

The trajectories observed in the data. Monthly averages are overlayed in orange.

```{r plot observed trajectories}
#generate plots of the observed KCCQOS trajectories over time for each subject
time_means <- subjects_frame_spline %>%
  group_by(time, treatment) %>%
  summarize(mean_observed = mean(observed), se_observed = sqrt(var(observed) / n())) %>%
  ungroup() %>%
  mutate(lower = mean_observed - 1.96*se_observed, upper = mean_observed + 1.96*se_observed)
  
plot_observed_trajectories <- ggplot(subjects_frame_spline, aes(x = time, y = 100*observed)) +
  geom_line(aes(group = subjects)) +
  geom_point(aes(group = subjects)) +
  geom_pointrange(data = time_means, aes(x = time, y = 100*mean_observed, ymin = 100*lower, ymax = 100*upper), size = 1.1, colour = "darkorange3") +
  geom_line(data = time_means, aes(x = time, y = 100*mean_observed), size = 1.0, colour = "darkorange3") +
  facet_grid(~ treatment) +
  labs(title = "Observed Trajectories", x = "Months", y = "Observed KCCQOS") +
  ylim(0,100) +
  scale_x_continuous(breaks = c(0,1,6,12), labels = c(0,1,6,12))
plot_observed_trajectories
```


# Individual Estimated KCCQOS
```{r plot spline trajectories}
#same as above but plot fitted trajectories
plot_spline_predictors <- ggplot(subjects_frame_spline, aes(x = time, y = 100*estimate, group = subjects)) +
  geom_line() +
  geom_point() +
  #ylim(0,1) +
  facet_grid(~ treatment) +
  labs(title = "Fitted Linear Trajectories", x = "Months", y = "Estimated KCCQOS") +
  ylim(0,100) +
  scale_x_continuous(breaks = c(0,1,6,12), labels = c(0,1,6,12))

plot_spline_predictors
```


```{r ppd check: observed versus fitted plot}
#Posterior predictive check
#baseline data is excluded from this plot 
#we basically asserted that it is similar between both groups which leads to very little variance in the baseline estimates and a strange sigmoidal artifact in the residuals
plot_fit <- ggplot(subjects_frame_spline %>% filter(time != 0), aes(x = 100*observed, y = 100*estimate, group = subjects)) +
  geom_point() +
  #ylim(0,1) +
  facet_grid( ~ treatment) +
  labs(title = "Fitted Linear Trajectories", x = "Observed KCCQOS", y = "Estimated KCCQOS") +
  ylim(0,100) +
  xlim(0,100)
plot_fit
```

```{r extract posterior draws}
#this chunk just gets draws from the fitted stan models

#joint draws
draws_joint <- extract(joint_fit_spline, pars = c("mu_theta_0", "mu_theta_1", "mu_theta_2", "gamma_1", "gamma_2", "beta_t", "beta_theta_1","beta_theta_2"))
draws_mu_theta_0_joint <- draws_joint$mu_theta_0 %>% as.numeric()
draws_mu_theta_1_joint <- draws_joint$mu_theta_1 %>% as.numeric()
draws_mu_theta_2_joint <- draws_joint$mu_theta_2 %>% as.numeric()
draws_gamma_1_joint <- draws_joint$gamma_1 %>% as.numeric()
draws_gamma_2_joint <- draws_joint$gamma_2 %>% as.numeric()
draws_beta_t_joint <- draws_joint$beta_t %>% as.numeric()
draws_beta_theta_1_joint <- draws_joint$beta_theta_1 %>% as.numeric()
draws_beta_theta_2_joint <- draws_joint$beta_theta_2 %>% as.numeric()

#joint draws with capped survival
draws_joint_capped <- extract(capped_joint_fit_spline, pars = c("mu_theta_0", "mu_theta_1", "mu_theta_2", "gamma_1", "gamma_2", "beta_t", "beta_theta_1","beta_theta_2"))
draws_mu_theta_0_joint_capped <- draws_joint_capped$mu_theta_0 %>% as.numeric()
draws_mu_theta_1_joint_capped <- draws_joint_capped$mu_theta_1 %>% as.numeric()
draws_mu_theta_2_joint_capped <- draws_joint_capped$mu_theta_2 %>% as.numeric()
draws_gamma_1_joint_capped <- draws_joint_capped$gamma_1 %>% as.numeric()
draws_gamma_2_joint_capped <- draws_joint_capped$gamma_2 %>% as.numeric()
draws_beta_t_joint_capped <- draws_joint_capped$beta_t %>% as.numeric()
draws_beta_theta_1_joint_capped <- draws_joint_capped$beta_theta_1 %>% as.numeric()
draws_beta_theta_2_joint_capped <- draws_joint_capped$beta_theta_2 %>% as.numeric()
#separate draws
draws_separate <- extract(separate_fit_spline_capped, pars = c("mu_theta_0", "mu_theta_1", "mu_theta_2", "gamma_1", "gamma_2","beta_t"))
draws_mu_theta_0_separate <- draws_separate$mu_theta_0 %>% as.numeric()
draws_mu_theta_1_separate <- draws_separate$mu_theta_1 %>% as.numeric()
draws_mu_theta_2_separate <- draws_separate$mu_theta_2 %>% as.numeric()
draws_gamma_1_separate <- draws_separate$gamma_1 %>% as.numeric()
draws_gamma_2_separate <- draws_separate$gamma_2 %>% as.numeric()
draws_beta_t_separate <- draws_separate$beta_t %>% as.numeric()
```


```{r distribution of thetas}
theta_labels <- c(theta_0 = "theta_0 (Baseline KCCQOS)", theta_1 = "theta_1 (1-Month KCCQOS)", theta_2 = "theta_2 (Post 1-Month Slope)")
qol_labels <- c("0" = "O obs", "1" = "1 obs", "2" = "2 obs", "3" = "3 obs", "4" = "4 obs")

has_qol_data <- c(rep(1, n_distinct(data_list_spline$Subjects)), rep(0, data_list_spline$I - n_distinct(data_list_spline$Subjects)))
table_qol_data <- c(table(data_list_spline$Subjects),  rep(0, data_list_spline$I - n_distinct(data_list_spline$Subjects))) 
thetas_1 <- pars_spline %>% 
  filter(grepl("theta_1\\[", term)) %>%
  separate(term, into = c("term","index"), sep = "\\[") %>%
  mutate(index = as.numeric(gsub("\\]","",index))) %>%
  mutate(qol_data = table_qol_data)
thetas_2 <- pars_spline %>% 
  filter(grepl("theta_2\\[", term)) %>%
  separate(term, into = c("term","index"), sep = "\\[") %>%
  mutate(index = as.numeric(gsub("\\]","",index))) %>%
  mutate(qol_data = table_qol_data)
thetas <- thetas_1 %>% bind_rows(thetas_2)
```


# Population Level KCCQOS Curves
Average observed KCCQOS values (dotted) and model population estimates (solid), with 95\% credible intervals.

```{r population level spline curves}
#actual curves
observed_population_curves <- qol_data %>%
  group_by(time, trialarm) %>%
  summarize(mean_KCCQOS = mean(KCCQOS, na.rm = T), se_KCCQOS = sqrt(var(KCCQOS, na.rm = T) / n())) %>%
  ungroup() %>%
  mutate(lower_KCCQOS = mean_KCCQOS - 1.96*se_KCCQOS, upper_KCCQOS = mean_KCCQOS + 1.96*se_KCCQOS) %>%
  mutate(treat_group = ifelse(trialarm ==  "TEST", "Treatment", "Control")) %>%
  mutate(estimation = "Observed") %>%
  select(time = time, mean = mean_KCCQOS, lower = lower_KCCQOS, upper = upper_KCCQOS, treat_group = treat_group, estimation = estimation)

#longitudinal growth curves from Reynolds 2011 paper. 
#time zero is the observed intercept
lgc_means <- c(.344, .362, .482, .615, .484, .692, .407, .667)
lgc_lower <- c(.313, .331, .441, .575, .438, .652, .350, .620)
lgc_upper <- c(.393, .376, .522, .654, .530, .733, .463, .713)

longitudinal_growth_curves <- data.frame('mean' = lgc_means, 'lower' = lgc_lower, 'upper' = lgc_upper) %>%
  mutate(time = observed_population_curves$time) %>%
  mutate(treat_group = observed_population_curves$treat_group) %>%
  mutate(estimation = "Original Longitudinal Growth Curves") %>%
  select(time, mean, lower, upper, treat_group, estimation)

#fitted curves from model
time_grid <- c(0,1,6,12)

curve_control_joint_capped <- pnorm(draws_mu_theta_0_joint_capped %*% t(as.numeric(time_grid == 0)) + draws_mu_theta_1_joint_capped %*% t(as.numeric(time_grid >= 1)) + draws_mu_theta_2_joint_capped %*% t(relu(time_grid-1))) %>% 
  melt(varnames = c("draws","time")) %>%
  group_by(time) %>%
  summarize(mean = mean(value), lower = quantile(value, .025), upper = quantile(value, .975)) %>%
  mutate(treat_group = "Control", time = time_grid, estimation = "Bayesian Joint Model (12-Month Survival)")

curve_treatment_joint_capped <- pnorm(draws_mu_theta_0_joint_capped %*% t(as.numeric(time_grid == 0)) + draws_mu_theta_1_joint_capped %*% t(as.numeric(time_grid >= 1)) + draws_gamma_1_joint_capped %*% t(as.numeric(time_grid >= 1)) + draws_gamma_2_joint_capped %*% t(relu(time_grid - 1)) + draws_mu_theta_2_joint_capped %*% t(relu(time_grid - 1))) %>% 
  melt(varnames = c("draws","time")) %>%
  group_by(time) %>%
  summarize(mean = mean(value), lower = quantile(value, .025), upper = quantile(value, .975)) %>%
  mutate(treat_group = "Treatment", time = time_grid, estimation = "Bayesian Joint Model (12-Month Survival)")

curve_control_joint <- pnorm(draws_mu_theta_0_joint %*% t(as.numeric(time_grid == 0)) + draws_mu_theta_1_joint %*% t(as.numeric(time_grid >= 1)) + draws_mu_theta_2_joint %*% t(relu(time_grid-1))) %>% 
  melt(varnames = c("draws","time")) %>%
  group_by(time) %>%
  summarize(mean = mean(value), lower = quantile(value, .025), upper = quantile(value, .975)) %>%
  mutate(treat_group = "Control", time = time_grid, estimation = "Bayesian Joint Model (30-Month Survival)")

curve_treatment_joint <- pnorm(draws_mu_theta_0_joint %*% t(as.numeric(time_grid == 0)) + draws_mu_theta_1_joint %*% t(as.numeric(time_grid >= 1)) + draws_gamma_1_joint %*% t(as.numeric(time_grid >= 1)) + draws_gamma_2_joint %*% t(relu(time_grid - 1)) + draws_mu_theta_2_joint %*% t(relu(time_grid - 1))) %>% 
  melt(varnames = c("draws","time")) %>%
  group_by(time) %>%
  summarize(mean = mean(value), lower = quantile(value, .025), upper = quantile(value, .975)) %>%
  mutate(treat_group = "Treatment", time = time_grid, estimation = "Bayesian Joint Model (30-Month Survival)")


fitted_population_qol_curves_joint <- bind_rows(curve_treatment_joint, curve_control_joint)
fitted_population_qol_curves_joint_capped <- bind_rows(curve_treatment_joint_capped, curve_control_joint_capped)
```


```{r separate model population level spline curves, include = FALSE}

#fitted curves from model
curve_control_separate <- pnorm(draws_mu_theta_0_separate %*% t(as.numeric(time_grid == 0)) + draws_mu_theta_1_separate %*% t(as.numeric(time_grid >= 1)) + draws_mu_theta_2_separate %*% t(relu(time_grid-1))) %>% 
  melt(varnames = c("draws","time")) %>%
  group_by(time) %>%
  summarize(mean = mean(value), lower = quantile(value, .025), upper = quantile(value, .975)) %>%
  mutate(treat_group = "Control", time = time_grid, estimation = "Bayesian Separate Model")

curve_treatment_separate <- pnorm(draws_mu_theta_0_separate %*% t(as.numeric(time_grid == 0)) + draws_mu_theta_1_separate %*% t(as.numeric(time_grid >= 1)) + draws_gamma_1_separate %*% t(as.numeric(time_grid >= 1)) + draws_gamma_2_separate %*% t(relu(time_grid - 1)) + draws_mu_theta_2_separate %*% t(relu(time_grid - 1))) %>% 
  melt(varnames = c("draws","time")) %>%
  group_by(time) %>%
  summarize(mean = mean(value), lower = quantile(value, .025), upper = quantile(value, .975)) %>%
  mutate(treat_group = "Treatment", time = time_grid, estimation = "Bayesian Separate Model")

fitted_population_qol_curves_separate <- bind_rows(curve_treatment_separate, curve_control_separate)
```

# Joint vs Separate Curves
```{r joint versus separate curves}
#put all estimates on the same plot in different panels
all_qol_curves <- bind_rows(longitudinal_growth_curves, fitted_population_qol_curves_joint, fitted_population_qol_curves_joint_capped, fitted_population_qol_curves_separate) %>%
  mutate(estimation = factor(estimation, levels = c("Original Longitudinal Growth Curves","Bayesian Separate Model","Bayesian Joint Model (12-Month Survival)", "Bayesian Joint Model (30-Month Survival)"))) %>%
  mutate(treat_group = ifelse(treat_group == "Control", "Standard Care", "TAVR"))

plot_all_qol_curves <- ggplot(all_qol_curves, aes(x = time, y = 100*mean)) +
  geom_ribbon(aes(fill = treat_group, ymax = 100*upper, ymin = 100*lower), alpha = .4) +
  geom_line(aes(color = treat_group)) +
  geom_point(aes(color = treat_group), size = 2.5) +
  xlab("Month") +
  ylab("Fitted Population Level KCCQ-OS") +
  facet_wrap(~ estimation) + 
  guides(fill = guide_legend(title="Group"), color = guide_legend(title = "Group")) +
  scale_x_continuous(breaks = c(0,1,6,12), labels = c(0,1,6,12)) +
  scale_y_continuous(limits = c(0,100), minor_breaks = seq(0,100,5), breaks = seq(0,100,10)) +
  theme_bw() +
  theme(panel.grid.major.y = element_line(colour = "grey40", size = .1, linetype = 'dashed'), panel.grid.major.x = element_blank(), panel.grid.minor = element_blank(), strip.text = element_text(size = 12, colour = "black"), axis.title = element_text(size = 14), axis.text = element_text(size = 10), legend.text = element_text(size = 12)) 
plot_all_qol_curves
```

# KCCQOS on the Same Plot with No SEs

```{r plot kccqos estimates tavr versus control}
#instead of separate panels for each method, overplot methods on the same panel (separated by treatment group).
#Do not show standard errors because they get too crowded

plot_qol_curves_trtgroup <- ggplot(all_qol_curves, aes(x = time, y = 100*mean)) +
  geom_line(aes(color = estimation), size = 1.5, alpha = .8) +
  geom_point(aes(color = estimation), size = 3) +
  xlab("Month") +
  ylab("Fitted Population Level KCCQ-OS") +
  facet_wrap(~ treat_group) +
  scale_x_continuous(breaks = c(0,1,6,12), labels = c(0,1,6,12)) +
  scale_y_continuous(limits = c(20,80), minor_breaks = seq(20,80,5), breaks = seq(20,80,10)) +
  theme_bw() +
  theme(panel.grid.major.y = element_line(colour = "grey40", size = .1, linetype = 'dashed'), panel.grid.major.x = element_blank(), panel.grid.minor = element_blank(), strip.text = element_text(size = 12, colour = "black"), axis.title = element_text(size = 14), axis.text = element_text(size = 10), legend.text = element_text(size = 12), legend.position = "bottom", legend.direction = "vertical", legend.title.align = .5)  + 
  guides(fill = guide_legend(title="Estimation Method", nrow = 4, byrow = TRUE), color = guide_legend(title = "Estimation Method"))

plot_qol_curves_trtgroup
```


# Results Summaries
```{r table 2}
#make a table of parameter estimates, there are some estimates that do not appear in the table 2 in the manuscript
#those estimates were referenced in the text

print_draw_results <- function(draws, decimals = 1){
  paste(round(mean(draws), decimals), " (", round(quantile(draws, .025), decimals), ", ", round(quantile(draws, .975), decimals), ")", sep = "")
}


########### spline results ###########

#1 month diff
month_1_qol_diff_joint_capped <- print_draw_results(100*(pnorm(draws_mu_theta_1_joint_capped + draws_gamma_1_joint_capped) - pnorm(draws_mu_theta_1_joint_capped)))

month_1_qol_diff_joint <- print_draw_results(100*(pnorm(draws_mu_theta_1_joint + draws_gamma_1_joint) - pnorm(draws_mu_theta_1_joint)))

month_1_qol_diff_separate <- print_draw_results(100*(pnorm(draws_mu_theta_1_separate + draws_gamma_1_separate) - pnorm(draws_mu_theta_1_separate)))

#6 month diff
month_6_qol_diff_joint_capped <- print_draw_results(100*(pnorm(draws_mu_theta_1_joint_capped + draws_gamma_1_joint_capped + 5*draws_mu_theta_2_joint_capped + 5*draws_gamma_2_joint_capped) - pnorm(draws_mu_theta_1_joint_capped + 5*draws_mu_theta_2_joint_capped)))

month_6_qol_diff_joint <- print_draw_results(100*(pnorm(draws_mu_theta_1_joint + draws_gamma_1_joint + 5*draws_mu_theta_2_joint + 5*draws_gamma_2_joint) - pnorm(draws_mu_theta_1_joint + 5*draws_mu_theta_2_joint)))

month_6_qol_diff_separate <- print_draw_results(100*(pnorm(draws_mu_theta_1_separate + draws_gamma_1_separate + 5*draws_mu_theta_2_separate + 5*draws_gamma_2_separate) - pnorm(draws_mu_theta_1_separate + 5*draws_mu_theta_2_separate)))

#12 month diff
month_12_qol_diff_joint_capped <- print_draw_results(100*(pnorm(draws_mu_theta_1_joint_capped + draws_gamma_1_joint_capped + 11*draws_mu_theta_2_joint_capped + 11*draws_gamma_2_joint_capped) - pnorm(draws_mu_theta_1_joint_capped + 11*draws_mu_theta_2_joint_capped)))

month_12_qol_diff_joint <- print_draw_results(100*(pnorm(draws_mu_theta_1_joint + draws_gamma_1_joint + 11*draws_mu_theta_2_joint + 11*draws_gamma_2_joint) - pnorm(draws_mu_theta_1_joint + 11*draws_mu_theta_2_joint)))

month_12_qol_diff_separate <- print_draw_results(100*(pnorm(draws_mu_theta_1_separate + draws_gamma_1_separate + 11*draws_mu_theta_2_separate + 11*draws_gamma_2_separate) - pnorm(draws_mu_theta_1_separate + 11*draws_mu_theta_2_separate)))

#survival hazards
treatment_hazard_joint_capped <- print_draw_results(exp(draws_beta_t_joint_capped), decimals = 2)
treatment_hazard_joint <- print_draw_results(exp(draws_beta_t_joint), decimals = 2)
treatment_hazard_separate <- print_draw_results(exp(draws_beta_t_separate), decimals = 2)


beta_theta_1_joint_capped <- print_draw_results(exp(draws_beta_theta_1_joint_capped*IQR(thetas_1$estimate)), decimals = 2)
beta_theta_2_joint_capped <- print_draw_results(exp(draws_beta_theta_2_joint_capped*IQR(thetas_2$estimate)), decimals = 2)

beta_theta_1_joint <- print_draw_results(exp(draws_beta_theta_1_joint*IQR(thetas_1$estimate)), decimals = 2)
beta_theta_2_joint <- print_draw_results(exp(draws_beta_theta_2_joint*IQR(thetas_2$estimate)), decimals = 2)

########### final results as table ############
results_table <- data.frame(
  "Posterior Quantity" = c("1-Month QOL Effect","6-Month QOL Effect", "12-Month QOL Effect", "Treatment Hazard", "Random Effect Hazard**", "Random Slope Hazard**"),
  "joint_capped Model (12 Months)" = c(month_1_qol_diff_joint_capped, month_6_qol_diff_joint_capped, month_12_qol_diff_joint_capped, treatment_hazard_joint_capped, beta_theta_1_joint_capped, beta_theta_2_joint_capped),
  "Joint Model (30 Months)" = c(month_1_qol_diff_joint, month_6_qol_diff_joint, month_12_qol_diff_joint, treatment_hazard_joint, beta_theta_1_joint, beta_theta_2_joint),
  "Separate Model" = c(month_1_qol_diff_separate, month_6_qol_diff_separate, month_12_qol_diff_separate, treatment_hazard_separate, NA, NA),
  "PARTNER Results" = c("13.3 (7.6, 19.0)", "20.8 (14.7, 27.0)", "26.0 (18.7, 33.3)", "0.55 (0.40, 0.74)*", "NA", "NA")
)
knitr::kable(results_table, caption = "QOL and survival treatment effect estimates for joint model and original PARTNER results. $^*$ = 1-year Kaplan-Meier estimate; $^{**}$ = hazard of 25th compared to 75th quantile of $\\theta_1$ or $\\theta_2$")
```


# References

- Ibrahim JG, Chu H, and Chen LM. Basic concepts and methods for joint models of longitudinal and surivval data. \textit{Journal of Clinical Oncology}. (2010).
- Gould L et al. Joint modeling of survival and longitudinal non-survival data: current methods and issues. Report of the DIA Bayesian joint modeling working group. \textit{Statistics in Medicine}. (2015).
- Leon M et al. Transcatheter aortic-valve implementation for aortic stenosis in patients who cannot undergo surgery. \textit{New England Journal of Medicine}. (2010).
- Reynolds MR et al. Health-related quality of life after transcatheter aortic valve replacement in inoperable patients with severe aortic stenosis. (2011).
````
