## Supplementary material for "Integrating Quality of Life and Survival Outcomes Cardiovascular Clinical Trials: Results from the PARTNER Trial": Code: read_partners_data.R

```
# read data 
library(haven)

#here we read in all data in the Created Data folder

all_files <- list.files("~/Desktop/Partner/Created Data/")
sas_files <- all_files[grep("sas7bdat", all_files)]
data_list <- list()
for(i in 1:length(sas_files)){
  data_list[[i]] <- read_sas(paste("~/Desktop/Partner/Created Data/", sas_files[i], sep = ""))
}
names(data_list) <- gsub("\\.sas7bdat", "", sas_files) #name files with their original name
```
