## Supplementary material for "Integrating Quality of Life and Survival Outcomes Cardiovascular Clinical Trials: Results from the PARTNER Trial": Code: README.txt

README for Joint Modeling of PARTNER Data

Paper: Integrating Quality of Life and Survival Outcomes Cardiovascular Clinical Trials: Results from the PARTNER Trial

Code author: Jacob Spertus

Paper Authors: Jacob Spertus, Laura Hatfield, David Cohen, Suzanne Arnold, Martin Ho, Phil Jones, Martin Leon, Bram Zuckerman, John Spertus

Files in this folder:

NOTE THAT DATA ARE NOT PUBLICLY AVAILABLE

read_partners_data.R - script to read in original data, called by joint_model_final.Rmd
joint_model_final.Rmd - rmarkdown script to run data analysis (some paths in first chunk will need to be changed to run)
separate_model_spline.stan - stan code to fit separate Bayesian survival and KCCQOS, called by joint_model_final.Rmd
joint_model_spline.stan - stan code to fit joint model, called by joint_model_final.Rmd
