## Appendix for "Integrating Quality of Life and Survival Outcomes Cardiovascular Clinical Trials: Results from the PARTNER Trial"

#### A Joint Model for KCCQ-OS and Survival

Here we describe a joint model for KCCQ-OS and survival. We use a Gaussian model for KCCQ-OS with mean estimated as a piecewise linear function of time and a Weibull model for survival with scale determined as a product of baseline hazard, treatment specific hazard, and random effect hazards.

Specifically, let $i \in \{1, ..., N\}$ indexes subjects and $j \in \{0, 1, 6, 12\}$ index time points. $Y_{ij}$ gives the KCCQ-OS score at time $j$ for subject $i$. $T_i$ is treatment for subject $i$, which does not depend on time. $S_i$ is survival time for subject $i$ and is possibly right censored. $I(j = k)$ is an indicator function which evaluates to 1 if $j = k$ (i.e. at time $k$) and 0 otherwise. The model is:

$$
\begin{aligned}
Y_{ij} = &(\mu_{\theta_0} + \theta_{0i}) \cdot I(j = 0) + && (1)\\
&(\mu_{\theta_1} + \gamma_1 T_i + \theta_{1i}) \cdot I(j \geq 1) + && (2)\\
&(\mu_{\theta_2} + \gamma_2 T_i + \theta_{2i}) \cdot \max(0, j - 1) + \epsilon_{ij} && (3)\\
S_i \sim &\text{Weibull}\{\alpha, \exp(-(\beta_0 + \beta_T \cdot T_i + \beta_{\theta_1} \cdot \theta_{1i} + \beta_{\theta_2} \cdot \theta_{2i}))\} && (4)
\end{aligned}
$$

The shared random effects appear in blue to highlight their presence in both the KCCQ-OS and survival models. Line (1) describes baseline KCCQ-OS: $\mu_{\theta_0}$ is the baseline mean across patients and $\theta_{0i}$ is a zero-centered random effect for patient $i$ giving there deviation from baseline. Line (2) describes 1-month KCCQ-OS: $\mu_{\theta_1}$ is the 1-month intercept for control subjects, $\gamma_1$ is the 1-month treatment effect, and $\theta_{1i}$ is the 1-month random effect for patient $i$. Finally, line (3) models a post 1-month KCCQ-OS slope: $\mu_{\theta_2}$ is the average slope in the control group, $\gamma_2$ is a treatment specific slope, and $\theta_{2i}$ is a random slope for patient $i$.

Specifying priors completes the model setup:

$$[\mu_{\theta_0}, \mu_{\theta_1}, \mu_{\theta_2}] \sim \mathcal{N}(0,1) \quad (5)$$
$$[\gamma_1, \gamma_2] \sim \mathcal{N}(0,1) \quad (6)$$
$$\epsilon_{ij} \sim \mathcal{N}(0,\sigma^2) \quad (7)$$
$$\theta_{0i} \sim \mathcal{N}(0,\sigma^2_{\theta_0}) \quad (8)$$
$$\theta_{1i} \sim \mathcal{N}(0,\sigma^2_{\theta_1}) \quad (9)$$
$$\theta_{2i} \sim \mathcal{N}(0,\sigma^2_{\theta_2}) \quad (10)$$
$$\alpha \sim \mathcal{N}^+(0,10) \quad (11)$$
$$[\beta_0, \beta_T, \beta_{\theta_1}, \beta_{\theta_2}] \sim \mathcal{N}(0,10) \quad (12)$$
$$[\sigma, \sigma_{\theta_0}, \sigma_{\theta_1}, \sigma_{\theta_2}] \sim t_3^+(0,1) \quad (13)$$

Lines (5) and (6) specify priors for the intercept and slope parameters in the longitudinal model. Recall that $Y_{ij}$ is approximately distributed as a standard normal after transformation such that most observations fall within [-2,2]. The $\mathcal{N}(0,1)$ priors on the intercept terms ($\mu_{\theta_0}$, $\mu_{\theta_1}$, and $\gamma_1$) place 95% prior mass on the interval [-2,2] and so are weakly informative. The $\mathcal{N}(0,1)$ priors are even less informative for the slope parameters since the time range (from 0 to 12) means that a slope of magnitude $|\mu_{\theta_2}| \geq 1$ (or $|\gamma_2| \geq 1$) is extremely unlikely. Similarly, the half-$t$ prior on $\sigma$ is also weakly informative because the range of $Y_{ij}$ seriously constrains the range of plausible values.

The priors on the random effects $\theta_{0i}$, $\theta_{1i}$, and $\theta_{2i}$ are specified hierarchically, such that their variances $\sigma_{\theta_0}$, $\sigma_{\theta_1}$, $\sigma_{\theta_2}$ are estimated from the data. Again considering the scale of $Y_{ij}$, the half-$t$ priors on the hierarchical variances are weakly informative.

In the survival model, the prior on the shape parameter $\alpha$ covers a large range of possible shapes. The priors on the linear coefficients in the hazard model provide a soft bound by placing 95% prior mass between [-20, 20]. If we were to exponentiate these to compute hazard ratios the range would be huge. Therefore the priors for the coefficients on the binary terms $\beta_0$ and $\beta_T$ are essentially non-informative.

The coefficients $\beta_{\theta_1}$ and $\beta_{\theta_2}$ control the degree of association between the health status and survival submodels. Because $\sigma_{\theta_1}$ and $\sigma_{\theta_2}$, the scales of $\theta_{1i}$ and $\theta_{2i}$, are not known in advance, priors for $\beta_{\theta_1}$ and $\beta_{\theta_2}$ are trickier to specify. For example, if $\sigma_{\theta_1}$ is small but $\theta_{1i}$ has a significant impact on survival then $\beta_{\theta_1}$ may be very large. One solution is to use a completely non-informative prior, but this can lead to an unstable posterior. We chose to use the $\mathcal{N}(0,10)$ prior, which allows a very large range for the hazards of $\beta_{\theta_1}$ and $\beta_{\theta_2}$. However, we recommend examining the posteriors for the $\beta_{\theta_1}$ and $\beta_{\theta_2}$ to ensure that the prior is not in conflict with the data. This is apparent if these coefficients end up being outside of the central mass of the prior, e.g. if the posterior mean for $\beta_{\theta_1}$ is -25. If this happens it may be a good idea to transform the data, for example by rescaling the time to years instead of months, which can help put $\theta_{2i}$ on a larger range.

### B Transformed KCCQ-OS Data

Figure S1 displays the sample quantiles of the KCCQ-OS data before and after transformation. If the data were perfectly normally distributed, the points would fall along the black line drawn in each plot. The trimming and probit transformation brings the data much closer to normality compared with the original data bounded between 0 and 1. There are still some deviations from normality in the tails and all KCCQ-OS scores of 100 are mapped to about 2.5 after transformation, leading to the "ledge" apparent in the right panel. There is no straightforward way to deal with the large number health status measurements exactly equal to 1, which is an artifact of the original measurement scale of the health status data.

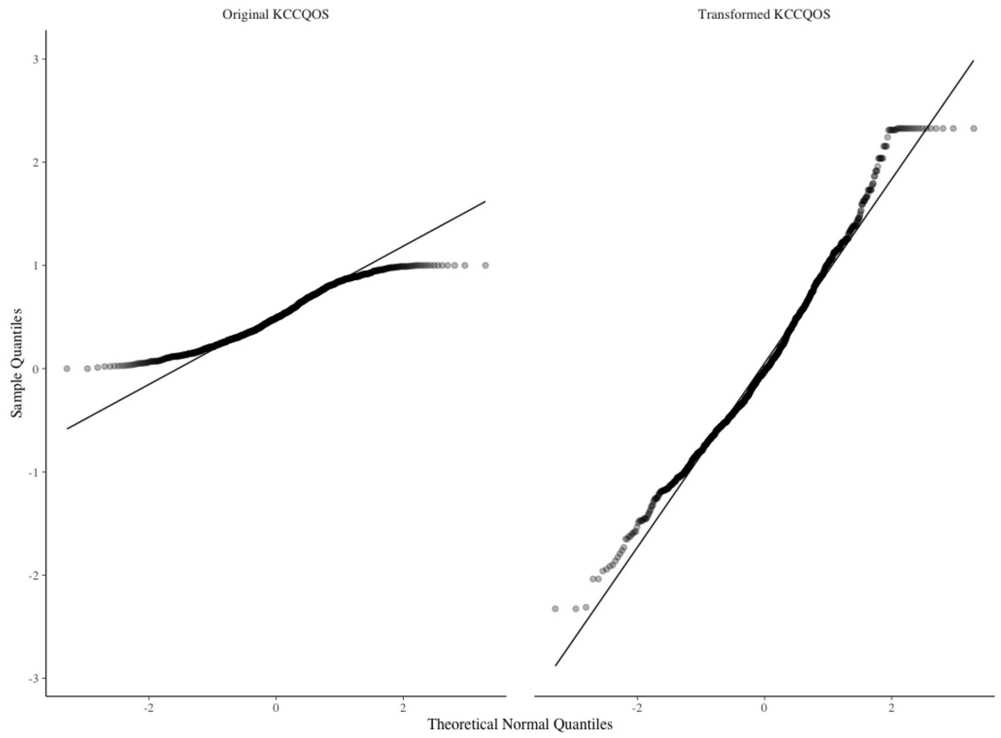


Figure S1: Plot of observed sample quantiles against theoretical normal quantiles.

### C Stan Code for Joint Model

```
//analysis of longitudinal data
data{
  // kccqos data
  int<lower=0> N; //number of total observations
  int<lower=0> I; //number of subjects
  vector[N] KCCQ; //quality of life for each patient at each time
  vector[N] Timepoint; //timepoints
  vector[I] Treatment; //treatment indicators
  int<lower = 1, upper = I> Subjects[N]; //indicator for subjects

  // survival data
  vector<lower=0>[I] Y; //survival time
  int Cen[I]; //censoring indicator
}

transformed data{
  // variables derived from Timepoint to make a piecewise linear function
  int<lower=0,upper=1> Time_0[N]; //intercept for baseline
  int<lower=0,upper=1> Time_1[N]; //value at 1-month and beyond
  vector[N] LaterTime; //(continuous) months beyond month 1
```

```
  for(n in 1:N){
    Time_0[n] = Timepoint[n] == 0;
    Time_1[n] = Timepoint[n] >= 1;
    LaterTime[n] = fmax(0, Timepoint[n] - 1);
  }
}

parameters{
  //qol parameters
  real mu_theta_0; //baseline qol
  real mu_theta_1; //1-month qol
  real mu_theta_2; //qol slope after 1-month

  real<lower = 0> sigma_theta_0; //standard deviation for baseline
  real<lower = 0> sigma_theta_1; //standard deviation for 1-month
  real<lower = 0> sigma_theta_2; //standard deviation for slopes-

  //real<lower = 0> sigma_theta_2; //standard deviation for slopes
  vector[I] theta_0_raw; //patient specific baseline
  vector[I] theta_1_raw; //patient specific 1-month
  vector[I] theta_2_raw; //patient specific post 1-month slope
  real gamma_1; //population level 1-month treatment effect
  real gamma_2; //population level treatment slope
  real<lower = 0> sigma_kccq; //standard deviation for kccq score

  //survival parameters
  real<lower=0> alpha; //shape parameter
  real beta_0; //intercept
  real beta_t; //treatment coefficient
  real beta_theta_1; //parameter to scale month 1 random intercepts
  real beta_theta_2;
}

transformed parameters{
  vector[N] linear_predictor; //fitted values
  vector[I] theta_0;
  vector[I] theta_1;
  vector[I] theta_2;
  theta_0 = theta_0_raw * sigma_theta_0; //theta_0 is scaled by sigma_theta_0 (its variance)
  theta_1 = theta_1_raw * sigma_theta_1; //theta_1 is scaled by sigma_theta_1 (its variance)
  theta_2 = theta_2_raw * sigma_theta_2;

  for(n in 1:N){
    linear_predictor[n] =
      (mu_theta_0 + theta_0[Subjects[n]]) * Time_0[n] +
      (mu_theta_1 + gamma_1 * Treatment[Subjects[n]] + theta_1[Subjects[n]]) * Time_1[n] +
      (mu_theta_2 + gamma_2 * Treatment[Subjects[n]] + theta_2[Subjects[n]]) * LaterTime[n];
  }
}
```

```
model{
  //longitudinal model priors
  theta_0_raw ~ normal(0, 1);
  theta_1_raw ~ normal(0, 1);
  theta_2_raw ~ normal(0, 1);
  sigma_theta_0 ~ student_t(3, 0, 1);
  sigma_theta_1 ~ student_t(3, 0, 1);
  sigma_theta_2 ~ student_t(3, 0, 1);
  sigma_kccq ~ student_t(3, 0, 1);
  mu_theta_0 ~ normal(0, 1);
  mu_theta_1 ~ normal(0, 1);
  mu_theta_2 ~ normal(0, 1);
  gamma_1 ~ normal(0, 1);
  gamma_2 ~ normal(0, 1);

  //survival model priors
  alpha ~ normal(0, 10);
  beta_0 ~ normal(0, 10);
  beta_t ~ normal(0, 10);
  beta_theta_1 ~ normal(0, 10);
  beta_theta_2 ~ normal(0, 10);

  //longitudinal model
  KCCQ ~ normal(linear_predictor, sigma_kccq);
  //survival model
  for(i in 1:I){
    if(Cen[i] == 0){
      Y[i] ~ weibull(alpha, exp(-(beta_0 + Treatment[i] * beta_t +
      beta_theta_1 * theta_1[i] + beta_theta_2 * theta_2[i]) / alpha));
  } else {
      target += weibull_lccdf(Y[i] | alpha, exp(-(beta_0 + Treatment[i] * beta_t +
       beta_theta_1 * theta_1[i] + beta_theta_2 * theta_2[i]) / alpha));
   }
 }
}

generated quantities{
  vector[N] bounded_predictor; //predictor on the original bounded scale
  bounded_predictor = Phi(linear_predictor);
}
```

### D Joint Parameters

Here we present results for joint parameter estimates from our joint model using 30-months of survival data. For the 30-month joint model, the hazard ratio of the 25th compared to the 75th percentile of random 1-month intercepts was 0.58 (95% CrI: 0.43, 0.76) and for the random slopes was 0.59 (95% CrI: 0.35, 0.91). In other words, a patient at the 75th percentile of the 1-month intercepts (which corresponds to a 1-month KCCQ-OS of 61 in the standard care group), is estimated to have about half the mortality risk as a patient at the 25th percentile (which corresponds to a 1-month KCCQ-OS of 36 in the standard care group). This implies that both the health status of patients at 1-month (after treatment) as well as the linear health status trajectories of patients after 1-month were linked to survival. We conclude that both random effects had a significant effect on survival as well as KCCQ-OS, and the model reflects the linkage between health status and survival.
